# Interferon-Induced PARP14-Mediated ADP-Ribosylation in p62 Bodies Requires an Active Ubiquitin-Proteasome System

**DOI:** 10.1101/2024.05.22.595402

**Authors:** Rameez Raja, Banhi Biswas, Rachy Abraham, Hongrui Liu, Che-Yuan Chang, Hien Vu, Anthony K. L. Leung

## Abstract

Biomolecular condensates are cellular compartments without enveloping membranes, enabling them to dynamically adjust their composition in response to environmental changes through post-translational modifications. A recent study has revealed that interferon-induced ADP-ribosylation (ADPr), which can be reversed by a SARS-CoV-2-encoded hydrolase, is enriched within a condensate. However, the identity of the condensate and responsible host ADP-ribosyltransferase remain elusive. Here, we demonstrate that interferon induces ADPr through transcriptional activation of PARP14, requiring both its physical presence and catalytic activity for condensate formation. Interferon-induced ADPr colocalizes with PARP14, and these PARP14/ADPr condensates contain key components of p62 bodies—including the selective autophagy receptor p62 and its binding partner NBR1, along with K48-linked and K63-linked polyubiquitin chains—but lack the autophagosome marker LC3B. Knockdown of p62 disrupts the formation of these ADPr condensates. Importantly, these structures are unaffected by autophagy inhibition but depend on both ubiquitin activation and proteasome activity. Taken together, these findings demonstrate that interferon triggers PARP14-mediated ADP-ribosylation in p62 bodies, which requires an active ubiquitin-proteasome system.

## INTRODUCTION

Biomolecular condensates are broadly defined as cellular compartments that are not enveloped by membranes^1–4^. Some, such as nucleoli, are constitutively present, while others, including DNA repair foci and stress granules, form in response to changes in cellular conditions. As components can freely diffuse in and out of these condensates, this constant exchange allows the condensates to rapidly adjust their composition over time, adapting to a changing environment. However, the mechanisms governing condensate composition remain a significant gap in the field.

Condensate formation can be regulated by post-translational modifications, such as ADP-ribosylation (ADPr)^5–8^—the addition of one or more ADP-ribose units onto proteins, regulated by a family of 17 human ADP-ribosyltransferases commonly known as PARPs. Within this family, four PARPs add poly(ADP-ribose) (PAR), 11 add mono(ADP-ribose) (MAR), and two are catalytically inactive (PARP9 and PARP13)^7,9^. PAR is critical for the structural integrity and function of DNA repair foci, stress granules, and nucleoli in human cells, while MAR is essential for forming *Sec* bodies under amino acid starvation in *Drosophila* cells^10,11^.

Recently, a novel class of ADPr-enriched condensates (hereafter ADPr condensates) was identified in the cytoplasm of human lung A549 epithelial cells during the search for coronavirus antivirals^12^. Notably, these condensates are induced by type I and II interferons (IFNα, β, and γ)^12^ and can be reversed by a SARS-CoV-2 macrodomain with hydrolase activity to remove MAR^13–16^. Within these condensates, ADPr regulation is mediated by the PARP9/DTX3L heterodimer^12^. However, as PARP9 is catalytically inactive as an ADP-ribosylation writer^9,17^, the specific enzyme responsible for MAR addition remains unidentified. This study delineates the identity of the interferon-induced PARP(s) involved in MAR addition, determines whether these ADPr condensates are part of a known class or a new cytoplasmic structure, and investigates the requirements for ADPr condensation.

Here, we report that these interferon-induced ADPr condensates are regulated by the MAR-adding ADP-ribosyltransferase PARP14, which shares the same genomic loci as PARP9 and DTX3L—all of which are transcriptionally activated by interferons^18^. Intriguingly, PARP14 functions as a dual-activity enzyme, also possessing a macrodomain with hydrolase activity similar to that observed in SARS-CoV-2^19–21^. Notably, ectopic expression of a PARP14 macrodomain mutant, deficient in ADP-ribosylhydrolase activity, leads to high levels of ADP-ribosylation and forms ADPr condensates, colocalizing with the mutant^19^. However, it was unclear if this occurs with the endogenous wild-type protein. This study demonstrates that PARP14 colocalizes with interferon-induced ADPr condensates, depending on its transferase activity.

In addition to PARP14’s transferase activity, we also identified that forming these ADPr condensates depends on p62, also known as sequestosome1 (SQSTM1)^22–24^. p62 serves as a central hub for signaling pathways and directs ubiquitinated proteins toward degradation via autophagy. Upon interacting with ubiquitinated proteins, p62 condenses *in vitro* and forms p62 bodies within cells^25,26^. These p62 bodies, heterogeneous in size, are present in a wide range of cell lines from various tissue origins, including both normal and cancerous types^24,27^. Diverse structures containing p62 and ubiquitin have been noted in liver cancers, as well as in various neurodegenerative diseases, such as Parkinson’s, Alzheimer’s, and Huntington’s disease^28–31^. Therefore, elucidating how alterations in the composition of p62 bodies affect their function in cell signaling or protein degradation could open novel therapeutic opportunities^32^.

In this work, we established that the ADPr-containing p62 bodies formed upon interferon treatment are distinct from the canonical ones. While these bodies contain ubiquitin^26,31,33^, they lack the autophagy marker LC3B and are not affected by autophagy inhibition; instead, they depend on an active ubiquitin-proteasome system. These compositional changes in p62 bodies likely reflect dynamic adaptations to the immune environment, facilitated in part by PARP14-mediated ADPr.

## RESULTS

### Interferon-Induced Cytoplasmic ADPr Condensates Depend on PARP14

To investigate the molecular mechanisms behind ADPr condensate formation, we initially examined their response to interferon. Among the three interferons tested, IFNγ was more potent than IFNα and β (Fig. S1A), hence chosen for further study. ADPr condensates formed as early as 10 h after IFNγ exposure (Fig. 1A). To assess if ADPr condensate formation depends on continuous or transient IFNγ exposure, cells were exposed to IFNγ for various durations before switching to IFNγ-free media for the remaining time of the 24-h observation period (Fig. 1B). Remarkably, just 1 h of IFNγ exposure was enough to induce ADPr condensate formation (Fig. 1B), indicating that continuous IFNγ presence is not necessary after initial exposure.

**Figure 1.**
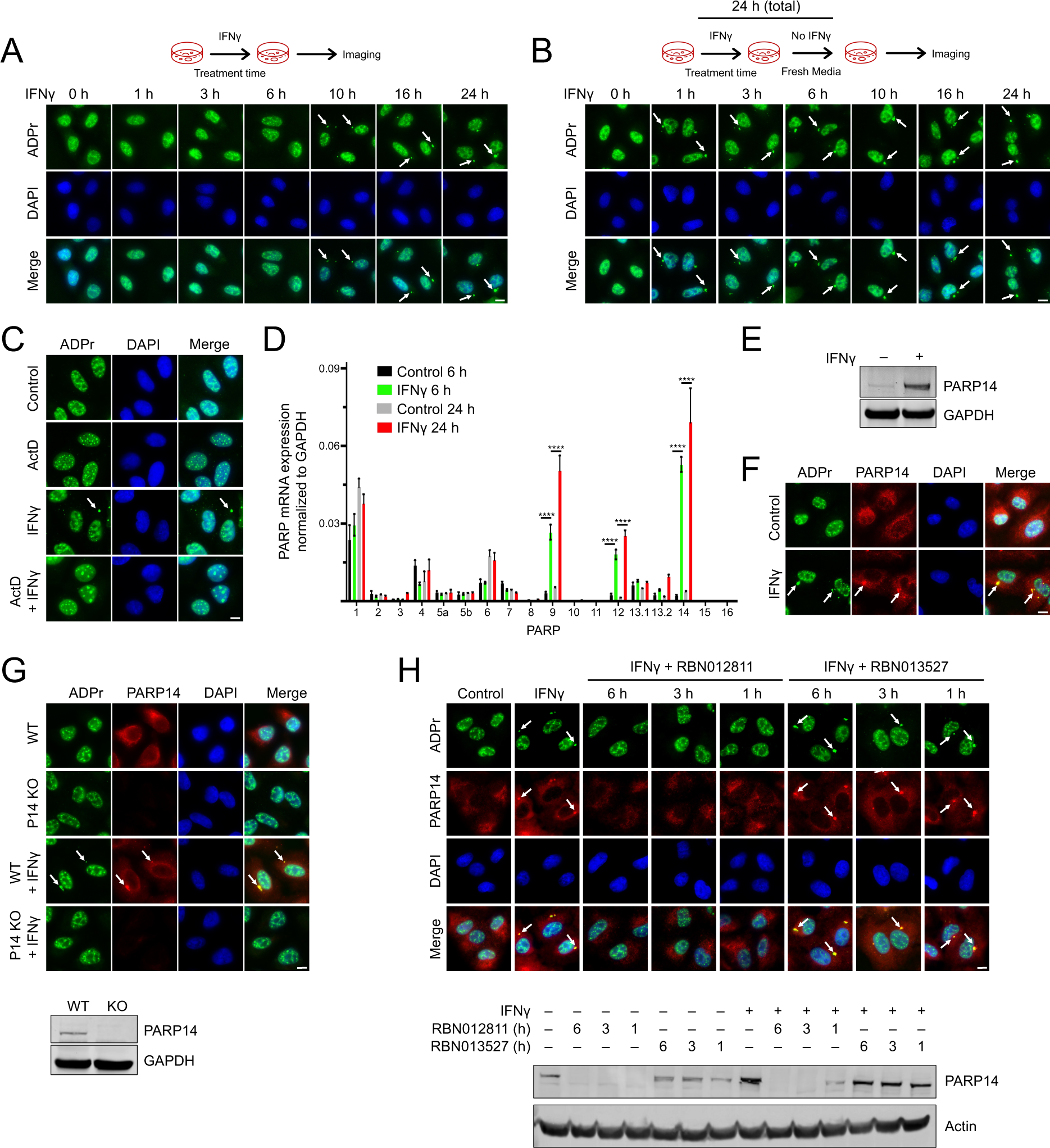
| Interferon-Induced Cytoplasmic ADPr Condensates Depend on PARP 14. (A) A549 cells were treated with IFNγ (500 IU/ml) for the indicated time points, stained with Pan-ADPr binding reagent MABE1016, and monitored for ADPr condensate formation. (B) ADPr condensate formation was monitored after cells were treated with IFNγ for the indicated time points and then replaced with fresh medium without IFNγ for a total duration of 24 h. (C) ADPr condensation formation was analyzed in cells pre-treated with Actinomycin D (ActD; 0.5 μg/ml) for 1 h followed by 24-h IFNγ treatment. (D) qPCR analyses of all human PARP expression in cells upon treatment with IFNγ for 6 h and 24 h. ****p < 0.0001, t-test, n = 3. (E) PARP14 protein levels were measured in cells treated with IFNγ for 24 h. (F) PARP14 and ADPr condensate colocalization was assessed in control and IFNγ-treated cells. (G) ADPr condensate formation was analyzed in A549 wild-type (WT) and PARP14 knockout (KO) cells treated with IFNγ for 24 h. The lower panel shows the corresponding PARP14 protein levels. (H) ADPr and PARP14 colocalization was analyzed in cells treated with IFNγ overnight, followed by treatment with either PROTAC RBN012811 (1 µM) targeting PARP14 or its negative control analog RBN013527 (1 µM) for the indicated time points. The lower panel shows the corresponding PARP14 protein levels. Scale bar, 10 µm.

The requirement for a 1-h IFNγ exposure and the ensuing 10-h delay before ADPr condensate formation indicates the possible involvement of a transcription program triggered by IFNγ. To determine whether ADPr condensate formation relies on transcription, we treated cells with the inhibitor Actinomycin D. Indeed, upon transcriptional inhibition, ADPr condensates were no longer observed (Fig. 1C).

As some PARPs are interferon-stimulated genes, we assessed the expression of all 17 human PARPs using qPCR after IFNγ treatment. PARP9, PARP12, and PARP14 were notably upregulated at 6 and 24 h post-treatment, with PARP14 showing the most significant increase (Fig. 1D and S1B). The significant upregulation of PARP14, a MAR-adding transferase, aligns with previous data indicating that the MAR-degrading SARS-CoV-2 Mac1 macrodomain can remove the ADPr signal within the condensates^13–16^. Western blot analyses demonstrated a corresponding induction of PARP14 protein post-IFNγ treatment (Fig. 1E). Immunostaining further revealed these ADPr condensates are also enriched with PARP14 (Fig. 1F). These results collectively highlight the critical role of IFNγ-induced transcription, particularly involving MAR-adding PARP14, in the formation of ADPr condensates.

To determine if the presence of PARP14 is crucial for ADPr condensate formation, we employed genetic depletion strategies, including siRNA, shRNA, and CRISPR/Cas9, to reduce PARP14 levels (Fig. 1G and S1C-H). In all cases, these IFNγ-induced ADPr condensates were no longer observed with PARP14 depletion (Fig. 1G and S1C-H).

Moreover, specific degradation of PARP14 using the proteolysis targeting chimera (PROTAC) inhibitor RBN012811 led to the disappearance of ADPr condensates within just 1 h (Fig. 1H)^34–36^. This effect was not observed with the negative control, RBN013527, an N-methylated analog^35^. Taken together, the physical presence of PARP14 is essential for ADPr condensate formation.

### PARP14 Catalytic Activity is Required for ADPr Condensate Formation and Co-Condensation with PARP14

As an orthogonal approach to determine which PARP activity is necessary for ADPr condensate formation, we screened a panel of PARP inhibitors (Fig. 2A and S2). This panel included inhibitors targeting different PAR-adding ADP-ribosyltransferases: Olaparib (PARPs 1/2), XAV939 (PARPs 1,2,5a/b), and MAR-adding ADP-ribosyltransferases: OUL-35 (PARP10), ITK6 (PARP10), ITK7 (PARPs 11/14), and RBN012579 (PARP14)^34,37–39^. Consistent with our investigations on transcriptional profiles and genetic ablation, only the inhibitors capable of inhibiting PARP14’s catalytic activity—specifically ITK7 and RBN012579 (hereafter RBN)^34,37^—suppressed ADPr condensate formation (Fig. 2A-B; S2). In alignment with the requirement for MAR-adding PARP14, these condensates were enriched with MAR (Fig. 2C). These findings underscore the crucial role of PARP14—both its physical presence and MARylation activity—in the formation of ADPr condensates.

**Figure 2.**
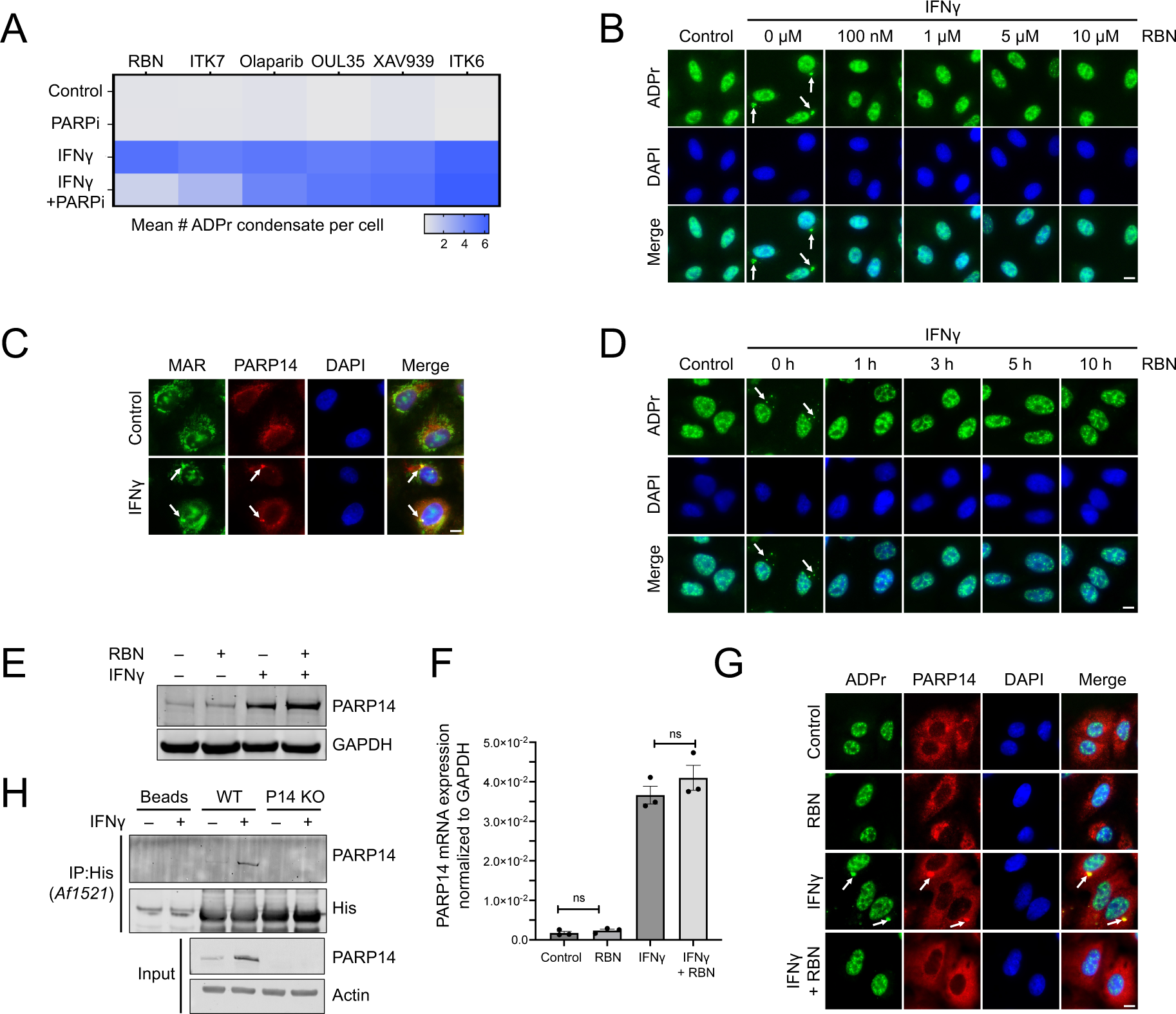
| PARP14 Catalytic Activity is Required for ADPr Condensate Formation and Co-Condensation with PARP14. (A) Heatmap showing the effect of different PARP inhibitors on ADPr condensate formation in A549 cells. (B) ADPr condensate formation was analyzed in cells pre-treated with different doses of RBN for 1 h, followed by 24-h IFNγ treatment. (C) MAR (HCA 355) and PARP14 colocalization was assessed after 24-h IFNγ treatment. (D) ADPr condensate formation was analyzed in cells treated with IFNγ overnight, followed by RBN treatment (10 μM) for the indicated time points. (E) PARP14 protein levels were measured in RBN-pretreated cells either alone or in the presence of IFNγ for 24 h. (F) PARP14 mRNA levels were measured after 6 h of IFNγ treatment either alone or in the presence of RBN by qPCR. ns = not significant, t-test, n = 3. (G) Colocalization of ADPr and PARP14 was assessed in control and RBN pretreated cells after 24-h IFNγ treatment. (H) A549 WT and PARP14 KO cells were treated with IFNγ for 24 h and subjected to immunoprecipitation assay using Pan-ADPr binding reagent (MABE1016) overnight. Ni-NTA resin was used to pull down His-tagged MABE1016, followed by western blot analyses. Scale bar, 10 µm.

Given RBN’s greater specificity and lower required dose for effective inhibition than ITK7 (Fig. 2B and S2G), we chose RBN for our in-depth PARP14 inhibition analyses. Notably, RBN significantly reduced the formation of ADPr condensates with as brief as 1-h treatment (Fig. 2D). Although RBN treatment resulted in elevated PARP14 protein levels without altering mRNA levels (Fig. 2E-F), PARP14 was no longer observed in its condensate form (Fig. 2G), indicating that the mere physical presence of PARP14 is not sufficient for its localization along with ADPr. Because PARP14 was pulled down by ADP-ribose binding Af1521 macrodomain following IFNγ treatment (Fig. 2H), PARP14 might be present in an ADP-ribosylated state within these condensates. Taken together, these observations suggest that the continuous presence of both ADPr and PARP14 in condensate form depends on the catalytic activity of PARP14.

### p62 is Required for Condensation of ADPr and PARP14 upon IFNγ Treatment

Next, we explored whether these cytoplasmic condensates, enriched with ADPr and PARP14 in an IFNγ-dependent manner, represent a known or novel cellular structure. We examined the colocalization of the IFNγ-induced ADPr signal with markers of various organelles (ER, Golgi, lysosome, mitochondria) and biomolecular condensates (stress granules, p-bodies), as well as macromolecular complexes such as immunoproteasome and proteasome (Fig. 3A and S3A). Among all structures tested, the ADPr signal did not colocalize with any markers, with the sole exception being the selective autophagy marker p62 (Fig. 3B and S3A). p62 can self-assemble into condensates *in vitro* and is crucial for forming “p62 bodies” within cells^25,26^. As expected, MAR and PARP14 were also detected in p62 bodies (Fig. 3B). Colocalization of p62, PARP14, and ADPr was also observed in the melanoma cell line A375 following IFNγ treatment, suggesting that this phenomenon is not restricted to lung cells (Fig. S3B). Given that p62 bodies exist constitutively in unstressed conditions^27^ (Fig. 3B), these findings suggest that PARP14 is localized to p62 bodies upon IFNγ treatment when ADP-ribosylation occurs.

**Figure 3.**
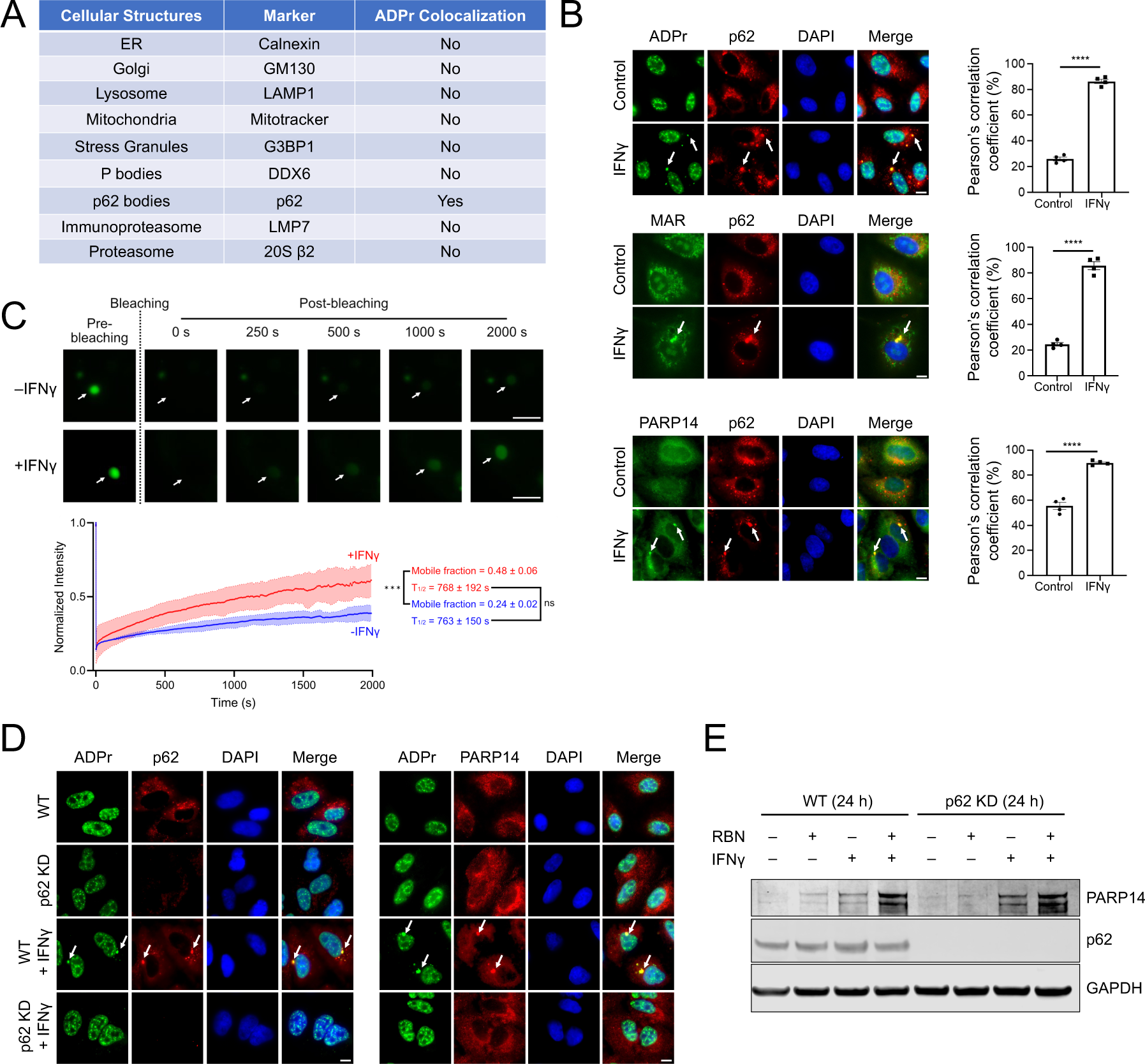
| p62 is Required for Condensation of ADPr and PARP14 upon IFNγ Treatment. (A) Table summarizing the colocalization of different cellular structure markers with ADPr condensates in A549 cells. (B) Colocalization of p62 with ADPr, MAR, and PARP14 was analyzed in cells after 24-h IFNγ treatment. Colocalization efficiency was quantified through Pearson’s correlation coefficient. ****p < 0.0001, t-test, n = 3. (C) FRAP analyses of A549 cells transiently transfected with GFP-p62 with or without 24-h IFNγ treatment. ***p < 0.001, t-test, n = 3. Scale bar, 5 µm. See the Methods section for the calculation of mobile fractions and T_½_. (D) ADPr and PARP14 condensate formation were analyzed in wild-type (WT) and p62 knockdown (KD) A549 cells after 24-h IFNγ treatment. (E) PARP14 and p62 protein levels in WT and p62 KD cells pretreated with RBN for 1 h followed by 24-h IFNγ treatment. Scale bar, 10 µm, unless otherwise stated.

Following IFNγ treatment, p62 bodies increased in the overall size distribution (Fig. S3C). Fluorescence Recovery After Photobleaching (FRAP) analyses of similarly sized p62 bodies revealed that IFNγ treatment increased the mobile fraction of p62 within the bodies. However, it did not alter the exchange dynamics between p62 bodies and the cytoplasm (Fig. 3C and S3D). Taken together, IFNγ treatment modulates the size, composition, and physical properties of p62 bodies.

To further investigate the role of p62 in ADPr condensates, we generated p62 knockdown clones from A549 cells using lentiviral transduction (Fig. S3E). p62 knockdown resulted in the loss of IFNγ-induced ADPr signal and PARP14 in the form of cytoplasmic condensates (Fig. 3D), despite PARP14 protein levels remaining unchanged in the presence of IFNγ (Fig. 3E). Therefore, p62 is required for the condensation of ADPr and PARP14 upon IFNγ treatment.

### PARP14-Mediated MARylation Facilitates p62 Co-Condensation

To investigate the interaction between p62 and PARP14, we immunoprecipitated p62 and probed for PARP14. p62 and PARP14 were associated with each other in untreated conditions, but more PARP14 was pulled down by p62 following treatment with IFNγ (Fig. 4A). As the protein and mRNA levels of p62 remained unchanged upon IFNγ treatment (Fig. 4A-B and S4A), the increased association is likely attributable to the upregulation of PARP14 expression (Fig. 4A). This association prompted us to inquire whether p62 could be a substrate of PARP14 for MARylation. We observed an increased MARylation of p62 upon IFNγ treatment; however, this increase was abrogated in cells lacking PARP14 (Fig. 4C), where p62 bodies were still present (Fig. 4D). Collectively, these data suggest that p62 undergoes increased MARylation by PARP14 upon IFNγ treatment.

**Figure 4.**
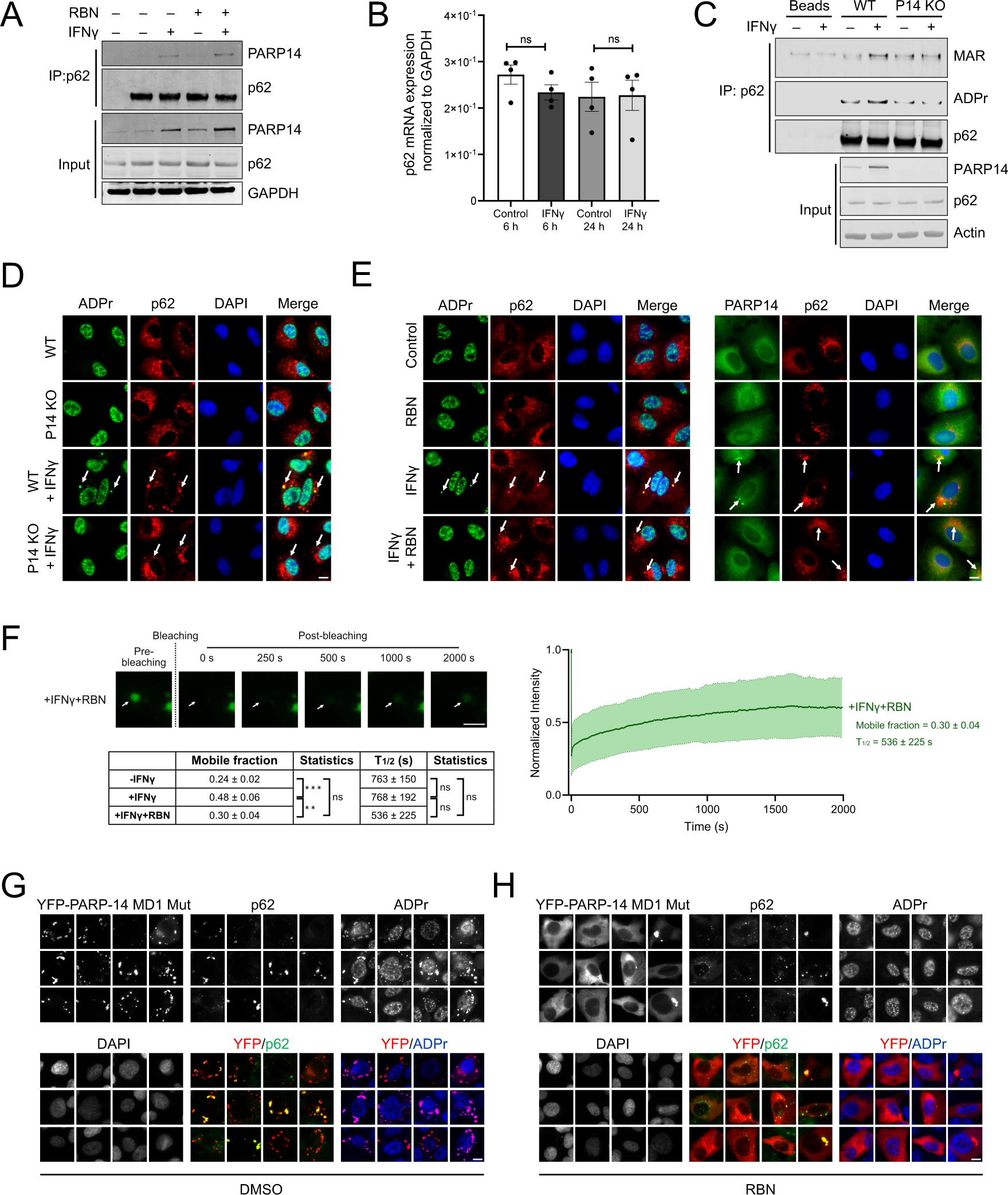
| PARP14-Mediated MARylation Facilitates p62 Co-Condensation. (A) A549 cells were treated with a different combination of IFNγ and RBN, and immunoprecipitates of p62 were probed for p62 and PARP14. (B) p62 mRNA levels were measured after 6 h and 24 h of IFNγ treatment by qPCR. ns = not significant, t-test, n = 4. (C) Immunoprecipitates of p62 were probed for MAR (HCA-355), ADPr and p62 from wild-type (WT) and PARP14 knockout (KO) A549 cells after 24-h IFNγ treatment. (D) ADPr and p62 colocalization was analyzed in WT or PARP14 KO cells after 24-h IFNγ treatment. (E) Colocalization of p62 with ADPr/PARP14 was analyzed in cells pretreated with RBN for 1 h before 24-h IFNγ treatment. (F) FRAP analyses of GFP-p62 transiently transfected in A549 cells followed by IFNγ treatment. 10 µM RBN was given for 6 h before FRAP analysis. Table summarizing FRAP analyses without IFNγ, with IFNγ, and with both IFNγ and RBN. ns, not significant; ***p < 0.01; *** p < 0.001; one-way ANOVA, n = 6. Scale bar, 5 µm. (G-H) U2OS cells were transiently transfected with YFP-PARP14 macrodomain 1 (MD1) mutant and treated with (G) DMSO control or (H) RBN (10 µM) for 6h. Scale bar, 10 µm, unless otherwise stated.

Importantly, upon treatment with the PARP14-specific inhibitor RBN, the association between PARP14 and p62 was not disrupted (Fig. 4A), indicating the association was not dependent on ADP-ribosylation. While the mRNA and protein levels of p62 remained unchanged with RBN treatment (Fig. 4A and S4A), the association was further increased, likely due to the increased PARP14 expression (Fig. 4A). Yet, the ADPr and PARP14 signals were no longer observed in p62 bodies (Fig. 4E and S4B), suggesting that the catalytic activity of PARP14, rather than its association with p62, is the prerequisite for ADPr condensation.

FRAP analyses further revealed that the mobile fraction of p62, in the presence of IFNγ and the PARP14 inhibitor RBN, was restored to levels comparable to those observed before IFNγ treatment, with no apparent change in recovery time (Fig. 4F and S3D). These findings suggest that PARP14, likely through its MAR-adding activity, alters the binding interactions within p62 bodies following IFNγ treatment.

To determine whether IFNγ is required for the co-condensation of PARP14-mediated ADPr and p62, we investigated the effects of a PARP14 mutant deficient in ADP-ribosylhydrolase activity (Fig. 4G). Previous studies demonstrated that transient transfection of this mutant into U2OS cells leads to the formation of cytoplasmic ADPr condensates colocalizing with PARP14, independent of IFNγ treatment^19^. Notably, a subset of condensates—particularly the larger ones—that contain both PARP14 and ADPr showed strong colocalization with p62 (Fig. 4G). Treatment with RBN under these conditions resulted in the disappearance of ADPr/PARP14 condensates while p62 bodies remained (Fig. 4H), further indicating that ADPr enriched in p62 bodies depends on the transferase activity of PARP14. These findings suggest that elevated PARP14-mediated ADP-ribosylation, whether induced by IFNγ or resulting from reduced hydrolase activity, can lead to its co-condensation with p62 bodies.

### ADPr-Enriched p62 Bodies Contain Ubiquitin but Lack Autophagy Marker LC3B

p62, a multifunctional scaffolding protein, comprises multiple domains that facilitate protein interactions critical for cell signaling and protein degradation (Fig. 5A)^22,24,25^. Inflammatory NF-κB signaling is mediated by the N-terminal self-associating PB1 domain, a ZZ-type zinc finger motif, and a TRAF6-binding domain (TB), whereas antioxidant signaling NRF2 activation is mediated through the Keap1-interacting region (KIR). These signaling domains are linked by an intrinsically disordered region that interacts with the autophagosome via LC3 (LIR). This LC3 interaction, coupled with the ubiquitin-associated (UBA) domain at p62’s C-terminus, targets ubiquitinated proteins for autophagy, positioning p62 as a selective autophagy receptor.

**Figure 5.**
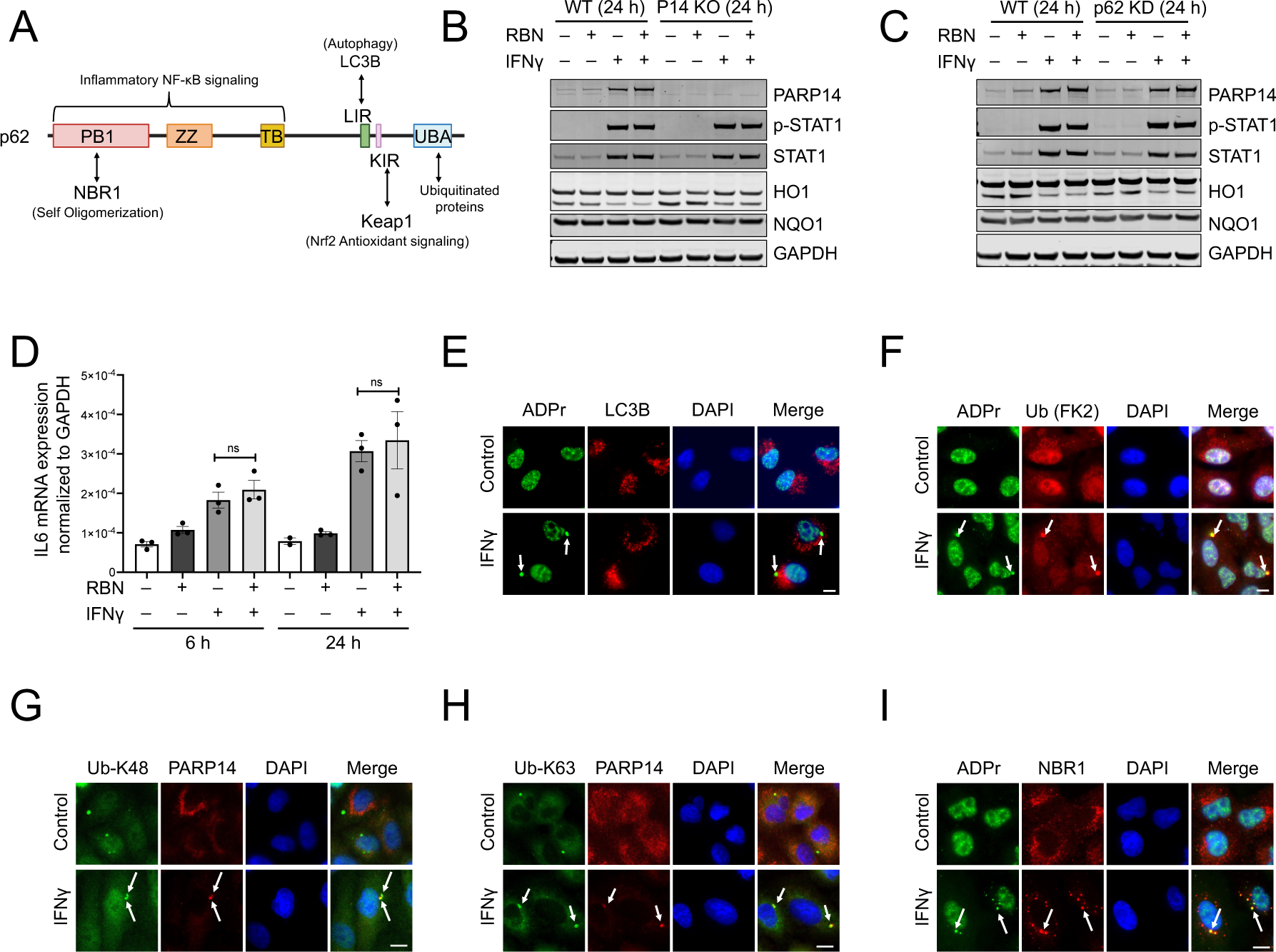
| ADPr-Enriched p62 Bodies Contain Ubiquitin but Lack Autophagy Marker LC3B. (A) Schematic of p62 domain architecture. (B-C) p-STAT1, STAT1, HO1, and NQO1 levels were compared in (B) A549 wild-type (WT) vs. PARP14 KO cells and (C) WT vs. p62 KD cells treated with different combinations of treatments with RBN and IFNγ. (D) IL-6 mRNA levels were measured in A549 cells treated with different combinations of RBN pretreatment for 1 h and IFNγ treatment for either 6 h or 24 h by qPCR. ns = not significant, t-test, n = 3. (E-I) A549 cells after 24-h IFNγ treatment were analyzed for colocalization between (E) ADPr and autophagy marker LC3B, (F) ADPr and ubiquitination (FK2), (G) PARP14 and Ub K48, (H) PARP14 and Ub K63, as well as (I) ADPr and NBR1. Scale bar, 10 µm.

We initially explored the activation of NF-κB and NRF2 signaling pathways by IFNγ and whether this activation is PARP14-dependent. Activation of the IFNγ signaling pathway involves the phosphorylation of STAT1 at tyrosine 701. STAT1 phosphorylation remained robust at 6 and 24 h in A549 PARP14 knockout cells or in wild-type parental lines treated with the PARP14 inhibitor RBN (Fig. 5B and S5A). This pattern was also observed in p62 knockdown cells (Fig. 5C and S5B), suggesting that the IFNγ signaling pathway is not dependent on p62 or PARP14.

No significant NRF2 signaling activation was observed with or without IFNγ treatment: its downstream targets, NQO1 and HO1, were not induced, regardless of PARP14 levels or activity, as evidenced respectively by genetic depletion and RBN inhibition (Fig. 5B and S5A). Similarly, NF-κB-induced genes, such as IL6 and OAS1, maintained consistent expression levels despite the presence of the PARP14 inhibitor (Fig. 5D and S5C). Taken together, these results suggest that under IFNγ stimulation, the regulation of NF-κB and NRF2 signaling activation in these cells does not depend on the physical presence and enzymatic activity of PARP14.

Next, we investigated the relationship between ADPr condensates and two major mechanisms of protein degradation mediated by p62. Normally, autophagy initiation is suppressed by the mTOR pathway. However, when inhibited with the mTOR inhibitor Torin-1, autophagy is induced, leading to increased autophagosome formation marked by LC3B on the membranes, which facilitates the recruitment of p62 and ubiquitinated proteins (Fig. S5D). However, in contrast to Torin-1-induced autophagy, the ADPr-containing p62 bodies induced by IFNγ treatment notably lacked LC3B (Fig. 5E).

Despite this difference, these p62 bodies that contain ADPr and PARP14 were enriched in ubiquitinated proteins (Fig. 5F-H and S5E-G), including both K63 and K48 linkages^25,26,31^, with the latter being known for its role in protein degradation. This colocalization with ubiquitin signals is similarly observed in canonical p62 bodies, which can be induced by puromycin (Fig. S5E-G), an amino acid analog that terminates protein synthesis and triggers polyubiquitination of these premature polypeptides^32,40^.

Additionally, we detected the presence of NBR1 (Fig. 5I)—a p62 homolog and binding partner—which not only promotes the formation of p62 bodies but also enhances the recruitment of ubiquitinated cargo cooperatively^26,33,41,42^. These data indicate that ADPr is enriched in p62 bodies that contain PARP14, NBR1, and ubiquitin but lack LC3B.

### Ubiquitin-Mediated Proteasome Activity is Required for PARP14 and ADPr Co-Condensation with p62 Bodies

The presence of ubiquitin, coupled with the absence of LC3B staining, suggested that autophagy might not be the primary pathway for ADPr regulation in these p62 bodies. This hypothesis was supported by data showing that treatment with autophagy inhibitors, such as chloroquine or Bafilomycin A1, did not impact the formation of ADPr condensates (Fig. 6A). Therefore, we decided to investigate proteasome pathway as an alternative mechanism of ADPr regulation.

**Figure 6.**
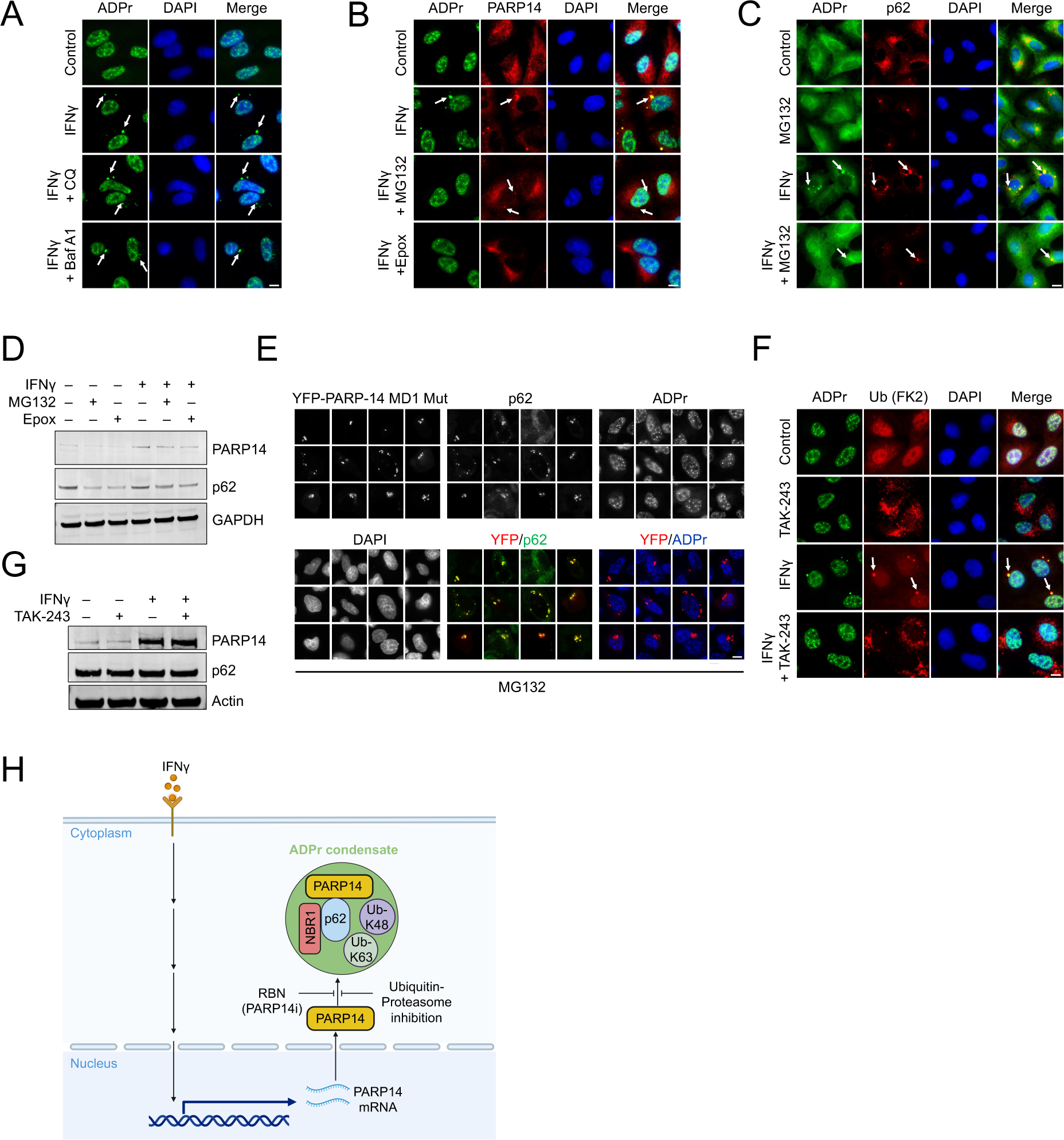
| Ubiquitin-Mediated Proteasome Activity is Required for PARP14 and ADPr Co-Condensation with p62 Bodies. (A) ADPr condensate formation was analyzed in A549 cells treated with IFNγ overnight, followed by the treatment with autophagy inhibitors 100 µM chloroquine (CQ) or 200 nM Bafilomycin A1 (Baf A1) for 6 h. (B) ADPr and PARP14 colocalization was assessed in cells treated with IFNγ overnight, followed by treatment with proteasome inhibitors 10 µM MG132 or 1 µM Epoxomicin (Epox) for 6 h. (C) ADPr and p62 colocalization was analyzed in cells treated with IFNγ overnight, followed by the treatment with 10 µM MG132 for 6 h (D) PARP14 and p62 protein levels were measured in cells treated with IFNγ overnight, followed by treatment with proteasome inhibitors 10 µM MG132 or 1 µM Epoxomicin for 6 h. (E) U2OS cells were transiently transfected with YFP-PARP14 macrodomain 1 (MD1) mutant and analyzed for p62, PARP14 (YFP), and ADPr colocalization after treatment of 10 µM MG132 for 6h. Nuclei were stained with DAPI. (F) ADPr and ubiquitination (FK2) colocalization was analyzed in cells treated with IFNγ overnight, followed by treatment with 1 µM E1 inhibitor TAK-243 for 6 h. (G) PARP14 and p62 protein levels were analyzed from samples in panel F. (H) Working model: IFNγ transcriptionally activates PARP14, leading to the formation of ADPr-containing p62 bodies, which require an active ubiquitin-proteasome system. Scale bar, 10 µm

To test whether active proteasome activity is required for IFNγ-induced ADPr in p62 bodies, we treated cells with proteasome inhibitor MG132. As expected, MG132 induced the formation of aggresomes, detected by the PROTEOSTAT® dye, which binds to misfolded proteins (Fig. S6A). However, the inhibition of the proteasome eliminated the ADPr condensate signals in a time-dependent manner (Fig. 6B and S6B-C). This phenomenon was further confirmed with two more selective proteasome inhibitors, Epoxomicin, and Bortezomib (Velcade; Fig. 6B and S6D).

Furthermore, upon proteasome inhibition, PARP14 signals within p62 bodies were significantly reduced (Fig. 6C and S6E-F), and the normally diffuse p62 signal became more prominent as condensates (Fig. 6C and S6F). Of note, inhibition of the proteasome slightly reduced p62 and PARP14 protein levels (Fig. 6D), which may partly explain the loss of ADPr and PARP14 signals.

The requirement of active proteasome was further demonstrated in U2OS cells transiently expressing PARP14 mutant deficient in ADP-ribosylhydrolase activity. Adding MG132 resulted in the loss of the ADPr signal, even when PARP14 and p62 were colocalized in condensates (Fig. 6E).

Proteasome activity can be mediated through both ubiquitin-dependent and ubiquitin-independent pathways^43–46^. To verify that the observed proteasome activity is indeed dependent on ubiquitination, we treated cells with TAK-243, a potent inhibitor of the ubiquitin-activating enzyme E1 (Fig. 6F). This blockade of ubiquitination initiation, similar to proteasome activity inhibition, resulted in the absence of ADPr condensates. However, unlike treatment with MG132, TAK-243 did not alter the levels of p62 and PARP14 (Fig. 6G). This indicates that changes in the levels of these two key proteins are not the primary factors regulating ADPr condensation.

Collectively, these data indicate that an active ubiquitin-proteasome system is required for the condensation of PARP14 and ADPr in p62 bodies.

## DISCUSSIONS

Here, we have identified that IFNγ activates a transcription program that increases PARP14 mRNA and protein levels, forming cytoplasmic ADPr condensates. Elevated levels of PARP14-mediated ADP-ribosylation facilitate its co-condensation with known components of p62 bodies, forming compositionally related structures that neither contain the autophagy marker LC3B nor are sensitive to autophagy inhibition. This class of IFNγ-induced ADPr condensates has the following characteristics:

### ADPr Condensates are Transient and Sensitive to PARP14 Levels and Activity

ADPr condensation requires both the physical presence and catalytic activity of PARP14. However, the mere presence is insufficient if its catalytic activity is inhibited. ADPr condensation is highly sensitive to both its levels and activity. Even brief interventions, such as 1 h catalytic inhibition or PROTAC-induced degradation of PARP14, can abolish ADPr condensation. This transient nature suggests that PARP14-mediated MARylation is continually removed by endogenous degraders, potentially including the macrodomain ADP-ribosylhydrolase activity inherent to this dual-activity enzyme^19–21^.

Notably, the expression of a PARP14 ADP-ribosylhydrolase-deficient mutant leads to elevated levels of ADP-ribosylation, with most substrates showing sensitivity to PARP14 transferase inhibition^19^. This observation underscores that many hydrolase targets are also transferase substrates, and the balance of their ADP-ribosylation is tightly regulated by the opposing enzymatic activities within PARP14. Our studies have identified two proteins, PARP14 itself and p62, that are ADP-ribosylated in a PARP14-dependent manner—both were also shown to be substrates for its hydrolase activity^19^. Since more ADP-ribosylation is observed for these and other substrates upon IFNγ treatment, cells may respond to the evolving immune environment by adjusting the enzymatic balance of PARP14—either via reducing hydrolase activity or enhancing transferase activity.

### High Levels of PARP14-Mediated ADP-Ribosylation are Sufficient to Induce their Condensation in a Specific Subset of p62 Bodies

PARP14 associates with p62 prior to IFNγ treatment, and these associations increase after treatment, likely due to the increased expression of PARP14. While this association is necessary, it is insufficient for co-condensation. PARP14 and p62 continue to associate when PARP14’s transferase activity is inhibited, yet they fail to co-condense, even with IFNγ treatment. Elevated protein levels typically increase condensation likelihood^1–4^. However, despite higher PARP14 levels after RBN treatment, PARP14 condensation does not occur without ADP-ribosylation. Instead, high levels of PARP14-mediated ADP-ribosylation are required for the co-condensation of PARP14, ADPr, and p62. This process can be triggered either by IFNγ treatment or by expressing an ADP-ribosylhydrolase-deficient PARP14 mutant without IFNγ treatment.

Such co-condensation occurs in p62 bodies with canonical components such as p62, its binding partner NBR1, and K48- and K63-linked polyubiquitin chains, yet lacking the autophagosome-associated LC3B. Our FRAP analyses further reveal that while the overall exchange rate of p62 with the cytoplasm remains constant, the fraction of p62 available for exchange increases. These compositional and dynamic differences align with the transient nature of these condensates, likely facilitating their response to changes in the immune environment.

### The Formation of ADPr-Containing p62 Bodies Requires an Active Ubiquitin-Proteasome System

Unlike canonical p62 bodies, these ADPr-containing ones are insensitive to autophagy inhibition. Instead, they are sensitive to the inhibition of ubiquitin activation and proteasome activity, indicating that their formation requires an intact ubiquitin-proteasome system. Crucially, p62 still forms condensates in the presence of proteasome or ubiquitin-activating enzyme E1 inhibitors; however, these condensates lack PARP14 and ADPr, highlighting a selective composition regulation within p62 bodies.

The requirement for an active ubiquitin-proteasome system led us to explore existing data on the crosstalk between ubiquitination and ADP-ribosylation. Recent data show that PARP9 and DTX3L bind to PARP14 and regulate its stability^47^. This stabilization, critical for increasing PARP14 levels, is independent of the proteasome^47^, implying it is unrelated to the IFN-induced ADPr condensate regulation.

Instead, the IFN-induced ADP-ribosylation depends on the ubiquitination of proteins that are destined for proteasome degradation, consistent with the enrichment of K48-linked polyubiquitin chains in these ADPr-containing p62 bodies. A potential enzyme responsible for this modification is the E3 ligase DTX3L, given its physical association with PARP14^47^. Consistent with this premise, knocking out DTX3L results in the loss of IFN-induced ADPr^12^. Our data now further suggest that not only the physical presence but also the enzymatic activity of DTX3L or other E3 ligases is critical, particularly their ability to ubiquitinate proteins with K48 linkages, which targets these poly-ubiquitinated proteins for proteasome degradation^46^. Identifying these ubiquitinated protein substrates destined for proteasome degradation during IFNγ treatment warrants further investigation.

It, however, remains unclear how inhibition of different components of the ubiquitin-proteasome system leads to decreased PARP14-mediated ADP-ribosylation in p62 bodies. While MG132 slightly reduces the amount of PARP14, TAK-243 does not have this effect. Thus, factors other than the modulation of PARP14 levels are likely contributing to the decreased ADP-ribosylation. One possibility is that the ubiquitin-proteasome system might reduce PARP14-mediated ADP-ribosylation by degrading either a repressor of PARP14 transferase activity or an activator of its hydrolase activity, following IFNγ treatment.

### Physiological and Pathological Implications

These IFNγ-induced ADPr condensates were discovered while developing a cell-based assay to screen inhibitors of viral ADP-ribosylhydrolase activity^12^. Expressing the first of the three macrodomains from SARS-CoV-2—the only one with hydrolase activity— reduces IFNγ-induced ADPr^12^. Remarkably, the three-macrodomain architecture of SARS-CoV-2 closely resembles that of PARP14, where only the first macrodomain exhibits ADP-ribosylhydrolase activity^19–21^. It would be intriguing to explore whether the virus usurps the host endogenous system, utilizing the same substrates and shifting the balance towards the hydrolysis of PARP14-mediated ADP-ribosylation in p62 bodies.

Besides its role in viral infection, IFNγ can also promote resistance to immunotherapy, such as those involving anti-PD-1 antibodies^36,48,49^. PARP14 upregulation has been observed in melanoma cell cultures derived from patient tumors resistant to immunotherapy with elevated IFNγ signaling^36^. Notably, in mouse models, restoring sensitivity to anti-PD-1 therapy can be achieved by PARP14 inhibitor RBN^36^. As ADPr-containing p62 bodies are also observed in melanoma cells following IFNγ treatment— and that such ADPr/PARP14 condensation is lost upon RBN treatment—understanding how PARP14-mediated ADP-ribosylation in p62 bodies functions could open new avenues for overcoming immunotherapy resistance.

## ACKNOWLEDGEMENT

We thank Drs. Phillip Sharp, Dani Cai, Anne Hamacher-Brady, Morgan Dasovich, as well as the Leung Lab members for critical comments on the manuscript. We thank Dr. Chenyao Wang and Ms. Sophia Cai for their assistance with image data analyses. We thank Drs. Christopher Sullivan and Anthony Fehr for sharing the A549 PARP14 KO cells and the parental lines, Dr. Vito Rebecca for A375 melanoma cells, Dr. Ivan Ahel for the YFP-PARP14 constructs, Dr. Anne Hamacher-Brady for the GFP-p62 construct, Dr. Michael Cohen for ITK6 and ITK7, Dr. Lari Lehtiö for OUL35, and Ribon Therapeutics, Inc. for various PARP14 inhibitors. We also thank Bluefield Innovation for the funding.

## AUTHOR CONTRIBUTIONS

Conceptualization: AKL and RR; Investigation and Validation: RR, BB, RA, HL; Resources: RR; Formal analysis: RR, BB, RA, HL, JC; Writing original draft: RR and AKL; Writing – review and editing: AKL, RR, RA, BB, HL, HV; Visualization: RR, BB, RA, HL, JC, AKL; Supervision: AKL, RR, RA; Project administration: AKL, RA; Funding acquisition: AKL

## MATERIALS AND METHODS

### Cell Lines and Reagents

A549 (CCL-185), HEK293T (CRL-3216) and U2OS (HTB-96) cells were purchased from the American Type Culture Collection (Manassas, VA, USA). A549 PARP14 KO^50^ and its wild-type counterpart were received as a kind gift by Dr. Christopher S. Sullivan, University of Texas, Austin. A375 Melanoma cell line was a kind gift from Dr. Vito W. Rebecca, Johns Hopkins University. A549 shPARP14 as well as shp62 knockdown cells were generated in the lab using lentiviral shRNA constructs TRCN0000290897 and TRCN0000007237 (Sigma). All cells were cultured in Dulbecco’s modified Eagle’s medium (DMEM; Gibco; Waltham, MA, USA) supplemented with 100 U/mL penicillin and streptomycin, 10% heat-inactivated fetal bovine serum (FBS; Gibco; Waltham, MA, USA) at 37 °C in 5% CO_2_ atmosphere. IFNγ (Sigma, #SRP3058) 500 IU/ml was used for most of the experiments at indicated time points in A549 cells. Table 1 details all drugs used along with IFNγ.

**Table 1:** List of Antibodies, Reagents and Chemicals Used.

| Antibodies | Source | Catalog Number | Host | Dilution WB | Dilution IF |
| --- | --- | --- | --- | --- | --- |
| SQSTM1/p62 (D5L7G) | CST | 88588 | Mouse | 1:1000 | 1:500 |
| SQSTM1/p62 (D10E10) | CST | 7695 | Rabbit | 1:1000 | 1:250 |
| PARP14 | Santa Cruz Biotechnology | sc-377150 | Mouse | 1:1000 | 1:250 |
| PARP14 | Ribon Therapeutics/<br>Genscript | U9980EK<br>130-3 | Rabbit |  | 1:250 |
| PARP14 | Abcam | Ab224352 | Rabbit |  | 1:100 |
| Pan-ADPr | Millipore | MABE101<br>6 | Rabbit | 1:500 | 1:500 |
| Mono-ADPr<br>(HCA-355) | Bio-Rad | abD33205 | Rabbit | 1:500 | 1:500 |
| Calnexin<br>(AF18) | Thermo Scientific | MA3-027 | Mouse |  | 1:250 |
| GM130 | BD Transduction<br>Laboratories | 610822 | Mouse |  | 1:250 |
| G3BP1-H10 | Santa Cruz<br>Biotechnology | SC-365338 | Mouse |  | 1:250 |
| LAMP1<br>(D401S) | CST | 15665 | Mouse |  | 1:100 |
| LC3B (E5Q2K) | CST | 83506 | Mouse |  | 1:500 |
| LMP7 (1A5) | CST | 13726 | Mouse |  | 1:50 |
| Stat1 | CST | 9172 | Rabbit | 1:1000 |  |
| Phospho-Stat1(58D6) | CST | 9167 | Rabbit | 1:1000 |  |
| HO1 (A-3) | Santa Cruz<br>Biotechnology | sc-136960 | Mouse | 1:1000 |  |
| NQO1 (A180) | Santa Cruz<br>Biotechnology | SC3187 | Mouse | 1:1000 |  |
| DDX6 | Sigma Aldrich | SAB4200<br>837 | Mouse |  | 1:100 |
| GAPDH (6C5) | EMD Millipore | MAB374 | Mouse | 1:10000 |  |
| β-Actin | Sigma Aldrich | A1978 | Mouse | 1:10000 |  |
| NBR1 (E6Q3F) | CST | 20145 | Rabbit |  | 1:100 |
| NBR1 (5C3) | Abcam | ab55474 | Mouse |  | 1:100 |
| Ub (P4D1) | Santa Cruz<br>Biotechnology | SC8017 | Mouse | 1:1000 | 1:50 |
| Ub (FK2) | EMD Millipore | 04-263 | Mouse |  | 1:500 |
| Ub-K48 | Sigma Aldrich | ZRB2150 | Rabbit |  | 1:100 |
| Ub-K63 | EMD Millipore | 05-1306 | Rabbit |  | 1:200 |
| GFP | Roche |  | Mouse | 1:1000 |  |
| 20S<br>Proteosome β2 | Santa Cruz<br>Biotechnology | SC-54810 | Mouse | | 1:50 |
| Cy5-Donkey<br>Anti-Mouse IgG | Jackson<br>ImmunoResearch | 715-175-150 | Donkey |  | 1:500 |
| Cy3-Donkey<br>Anti-Mouse IgG | Jackson<br>ImmunoResearch | 715-165-150 | Donkey |  | 1:500 |
| FITC-Fab<br>Donkey Anti-Rabbit IgG | Jackson<br>ImmunoResearch | 711-097-003 | Donkey |  | 1:500 |
| Cy5-Donkey<br>Anti-Rabbit IgG<br>(H+L) | Jackson<br>ImmunoResearch | 711-175-152 | Donkey |  | 1:500 |
| FITC-Donkey<br>Anti Mouse IgG<br>(H+L) | Jackson<br>ImmunoResearch | 715-095-150 | Donkey |  | 1:500 |
| **Reagents/Chemicals** |  |  |  |  |  |
| MG132 | Sigma Aldrich | C2211 |  |  |  |
| Epoxomicin | MedChem Express | HY-13821 |  |  |  |
| Bortezomib | MedChem Express | HY-10227 |  |  |  |
| Torin1 | MedChem Express | HY-13003 |  |  |  |
| Rapamycin | MedChem Express | HY-10219 |  |  |  |
| Chloroquine | CST | 14774 |  |  |  |
| BafilomycinA1 | MedChem Express | HY-100558 |  |  |  |
| ITK7 | Sigma Aldrich | SML2669 |  |  |  |
| Olaparib | Selleckchem | AZD2281 |  |  |  |
| XAV-939 | Selleckchem | S1180 |  |  |  |
| RBN012759 | MedChem Express | HY-136979 |  |  |  |
| RBN012811 | Ribon therapeutics | N/A |  |  |  |
| RBN013527 | Ribon therapeutics | N/A |  |  |  |
| OUL35 | Dr. Lari Lehtio |  |  |  |  |
| ITK6 | Dr. Michael Cohen |  |  |  |  |
| Actinomycin D | CST | 15021 |  |  |  |
| PROTEOSTAT<br>Aggresome<br>Detection kit | Enzo Life Sciences | ENZ-51035 |  |  |  |
| Puromycin | Invivogen | ant-pr-1 |  |  |  |
| IFNγ human | Sigma Aldrich | SRP3058 | | | |
| Lipofectamine 2000 | Invitrogen | 11668019 |  |  |  |

### Lentivirus Transduction and Stable Cell Line Generation

HEK-293T cells were seeded into a 10 cm dish to reach approximately 80% confluence by the following day. 16 h after seeding, cells were transfected using Lipofectamine 2000 (Invitrogen, 11668019) according to the manufacturer’s instructions with 18 µg of the following plasmid cocktail: lentiviral shRNA constructs from Sigma Aldrich (shPARP14 TRCN0000290897 and TRCN0000296754 or shp62 TRCN0000007237 and TRCN0000007236), psPAX2 (Addgene, #12260), and pMD2.G (Addgene, #12259) in a ratio of 3:2:1, respectively, to generate lentiviral particles. 48 h post-transfection, the supernatant was collected, centrifuged at 500 × g, and filtered through a 0.45-μm low-protein-binding filter. The resulting virus suspension containing lentiviral particles was used to transduce A549 cells overnight. Fresh media containing puromycin (Sigma) at a concentration of 1 μg/ml was then added for selection, and single-cell clones were picked for shRNA-based genetic depletion of PARP14 and p62 in A549 cells.

### Quantitative Real-time PCR

Total RNA was isolated from A549 cells under different treatment conditions using the High Pure RNA Isolation Kit (Roche), following the manufacturer’s instructions. The RNA was quantified and reverse transcribed with random hexamers, using the SuperScript VILO cDNA Synthesis Kit (Invitrogen) at 25°C for 10 min, 50°C for 10 min, and 85°C for 5 min. The resulting cDNA was mixed with the appropriate primers (Integrated DNA Technologies; Coralville, IA) and SYBR Green PCR Master Mix (Applied Biosystems) and then amplified for 40 cycles (30 s at 94°C, 40 s at 60°C, and 30 s at 72°C) on a 7500 Fast Real-Time PCR System (Applied Biosystems). The target gene expression was normalized to GAPDH, and the relative gene expression level was calculated using the 2^-ΔCT method. All primer sequences are listed in Table 2.

**Table 2:**
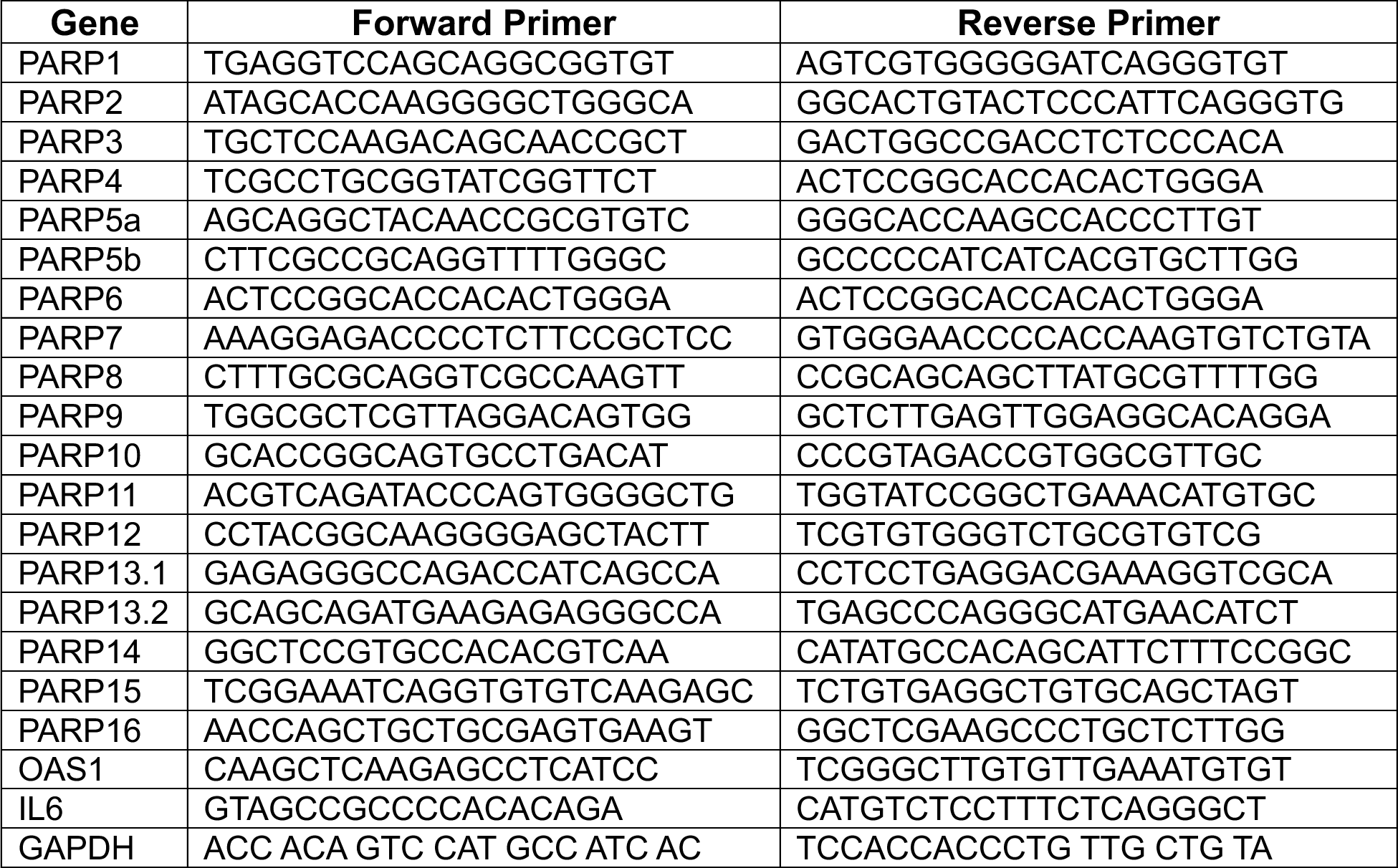
List of Primers Used.

### Immunoblotting

Cells were lysed in RIPA lysis buffer (CST, #9806) in presence of protease (Roche, # 11873580001) and phosphatase (Roche, #4906845001) inhibitors, 10 μM Olaparib and 10 μM PDD (PARG inhibitor) at 4^°^C for 30 min and centrifuged at 21,130 g at 4^°^C for 20 minutes. The lysates were quantitated using BCA assay (Thermo Scientific, #23225) and 20μg of proteins were electrophoresed on SDS–PAGE gels and then transferred to PVDF membrane (Thermo Scientific, # 88518). The membranes were blocked using 5% Bovine Serum Albumin (BSA) (Sigma, #A9647) in 1X TBST buffer (150 mM NaCl; Tris, pH 7.4; 0.1% Tween-20) for 1 h at room temperature. Blots were probed overnight at 4°C with primary antibodies, washed with TBST (3 times, 10 min each), and incubated with the appropriate secondary antibodies for 1 h at room temperature. Blots were washed again with TBST (3 times, 10 min each), and signals were detected on the Odyssey CLx imaging system using Image Studio Lite software (LI-COR). Western blot image data was processed by ImageJ software. The primary and secondary antibodies used are available in Table 1.

### Immunoprecipitation

Cells were lysed in immunoprecipitation buffer containing 20 mM HEPES (pH 7.5), 150 mM NaCl, 10 mM NaF, 1.5 mM MgCl_2_, 10 mM β -glycerophosphate, 2 mM EDTA, 5 mM Na-pyrophosphate, 1 mM Na_3_VO_4_, 1% (v/v) Triton X-100, 0.2% NP-40, and protease and phosphatase inhibitors and 10 μM Olaparib and 10 μM PDD (PARG inhibitor) at 4^°^C for 30 min and centrifuged at 21,130 g at 4^°^C for 20 min. The supernatant was incubated overnight with antibodies at 4^°^C, followed by incubation with Protein A/G PLUS-Agarose beads (Santa Cruz Biotechnology, #SC-2003) or Ni-NTA resin (Thermo Scientific, # 88222) for 2-4 h with rotation. The related beads were washed with IP buffer and boiled in 2x LDS buffer (Novex) for 10 min at 70^°^C. For Ni-NTA pull down, imidazole was added to 2x LDS buffer at 300mM concentration to facilitate the release of immunoprecipitated protein in 2x LDS buffer. The samples were run in SDS–PAGE gel and transferred onto PVDF membranes (Bio-Rad) and processed for western blot analysis.

### Transient Transfections

U2OS cells (5 × 10^4^ cells per well) were transiently transfected with 300 ng of plasmid construct YFP-PARP14 MD1 Mut (Kind gift from Dr. Ivan Ahel, University of Oxford) using Lipofectamine 3000 (Invitrogen, L3000001) as per manufacturer’s instructions. The media was replaced 6 h post-transfection. Cells were fixed with 4% PFA 24 h post-transfection. If inhibitors like MG132 (10 µM) and RBN (10 µM) were used, they were added 24 h post-transfection, with a treatment time of 6 h followed by cell fixation.

### Immunofluorescence

A549 cells upon IFNγ and/or drug treatment (5 × 10^4^ cells per well) or U2OS cells transiently transfected and/or drug treatment (5 × 10^4^ cells per well) in Ibidi µ-24 well plate were fixed with 4% paraformaldehyde (PFA) for 20 min. Following fixation, cells were permeabilized using 0.2% Triton X-100 in PBS for 20 min, washed twice with PBS, and blocked with 5% BSA in PBS for 1 h at room temperature and then labeled overnight with corresponding primary antibodies. The next day, cells were washed three times with PBS, and secondary antibodies (Jackson Immunoresearch) were added. Nuclei was stained with DAPI (5 µg/ml) and washed thrice with PBS before imaging. Images were acquired using a Leica Thunder Imaging System with 20X or 40X magnification and processed with Leica LAS X software and Image J.

Colocalization data was analyzed on ImageJ software using Pearson’s Correlation Coefficient images were analyzed with Fiji (ImageJ). Initially, the color channels of the images were separated. Nonspecific signals were minimized by applying noise median and despeckling tools to correct the background. After these steps, the Colocalization plugin was used to analyze how much two signals overlap. p62 bodies were identified with the built-in “Analyze Particles…” command with particle size set at greater than 0.5µm^2^ and particle circularity greater than 0.7 and plotted.

### Fluorescence Recovery After Photobleaching (FRAP) Analysis

FRAP experiments were performed on a Leica Thunder Imaging system equipped with an infinity scanner for photobleaching and a Leica DFC9000 GTC camera for fluorescence imaging. A549 cells (5 × 10^4^ cells per well) were transiently transfected with 500 ng GFP-p62 using Lipofectamine 2000 (Invitrogen,11668019) according to the manufacturer’s instructions. 6 h after transfection, the cells were treated with IFNγ overnight, followed by RBN treatment for 6 h, before performing FRAP analysis. GFP-tagged p62 condensates were photobleached using 100% 488 nm laser irradiation for 50 iterations. Time-lapse images were acquired post-bleaching every 10 seconds to monitor fluorescence recovery using a 63x oil objective (HC PL APO 63x/1.40-0.60 OIL), and fluorescence intensity within the bleached ROI and a reference unbleached region was measured using ImageJ. The fluorescence intensities were normalized by calculating their ratios to the fluorescence intensity of the same ROI before photobleaching. The recovery curves were fit by a single exponential equation:

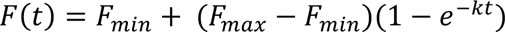

Where:

- *F*(*t*) is the normalized fluorescence intensity at time *t*,
- *F_max_* is the maximum fluorescence intensity recovered,
- *F_min_* is the normalized fluorescence intensity right after photobleaching,
- *k* is the rate constant.

The mobile fraction and half-time of recovery (T_1/2_) were calculated as:

- Mobile Fraction = *F_max_* − *F_min_*
- T_1/2_ is time *t* when the normalized fluorescence intensity equals to 50% (i.e., 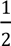 (*F_max_* − *F_min_*))

### Statistical Analysis

All statistical analyses were performed using GraphPad Prism 10 software.

For qPCR analysis error bars represent the standard deviation data from three independent experiments. The differences in statistical significance between the two groups were tested via a two-tailed t-test, and multiple groups were tested via one- or two-way ANOVA test. All values are means of SEM of the indicated independent experiments. ns> 0.05; *P < 0.05; **P < 0.01, ***P < 0.001, ****P < 0.0001.

For FRAP experiments, one-way ANOVA was performed to compare different experimental conditions. Controls, including unbleached regions within the same sample, were used to assess photobleaching and phototoxicity effects, and experiments were repeated in multiple independent trials to ensure reproducibility.

For Western blot, co-IP, and confocal experiments, unless indicated otherwise, results are representative of at least three independent experiments.

