## Supplementary Figures for "Interferon-Induced PARP14-Mediated ADP-Ribosylation in p62 Bodies Requires an Active Ubiquitin-Proteasome System"

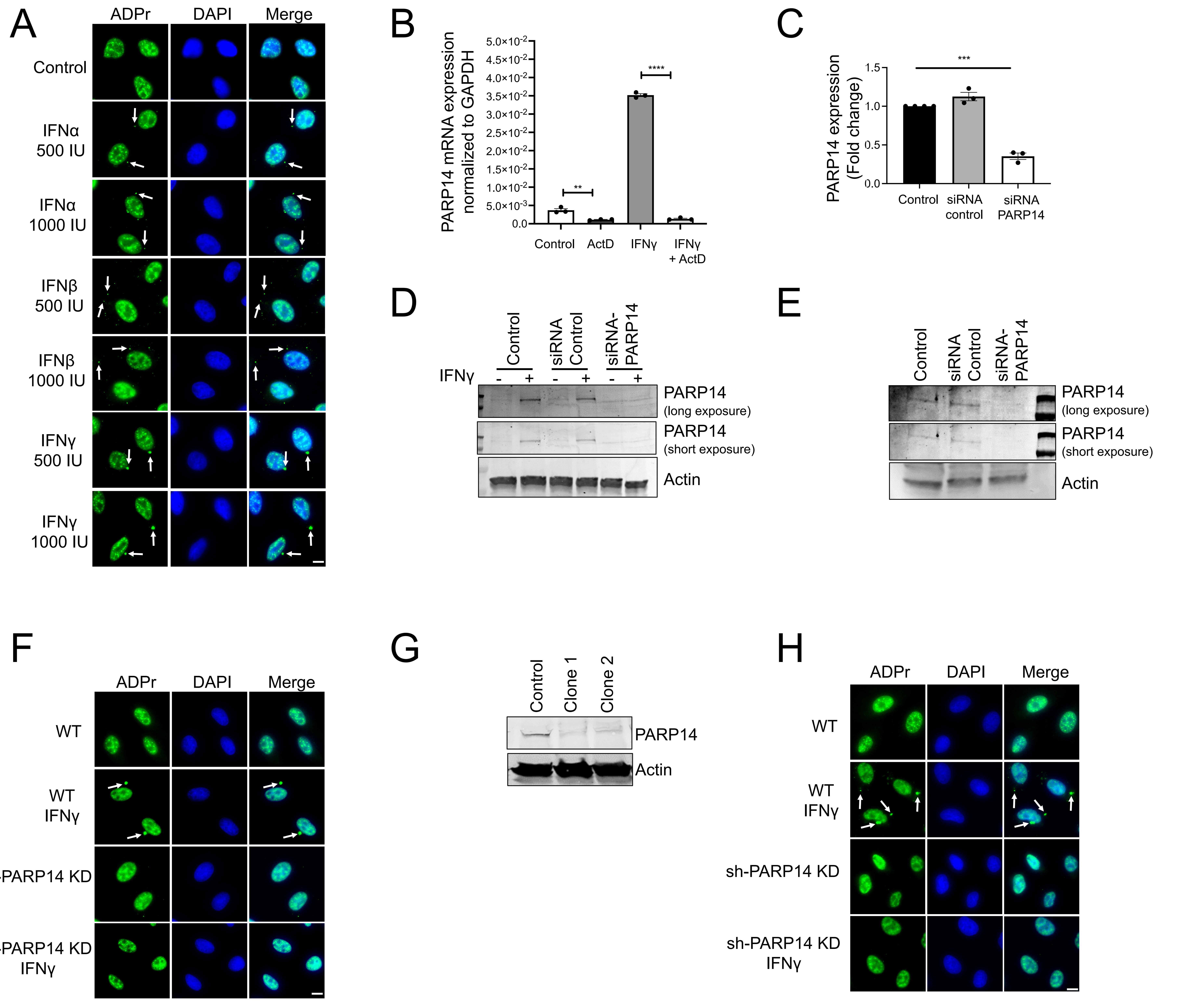


**Figure S1**

(A) ADPr condensates were monitored after cells treated with IFNα, β, γ (500 IU/ml). (B) PARP14 mRNA was measured by qPCR in A549 cells pretreated with Actinomycin D (ActD 0.5μg/ml), followed by IFNγ treatment of 6 h. **$p < 0.01$; ****$p < 0.0001$, t-test, n = 3. (C) PARP14 mRNA was measured by qPCR after 48 h of siRNA transfection. ***$p < 0.001$, t-test, n = 3. (D) PARP14 protein levels were measured in A549 cells after 48 h of siPARP14 transfection. (E) PARP14 protein levels were assessed in cells transfected with siPARP14 or control siRNA after IFNγ treatment. (F) ADPr condensates were analyzed in either wild-type or siPARP14 transfected cells with or without IFNγ treatment. (G) PARP14 protein levels were measured in A549 stably transduced with shRNA against PARP14 using lentivirus. (H) ADPr condensates were analyzed in either wild-type or shPARP14 knockdown cells after 24-h IFNγ treatment. Scale bar, 10 μm.

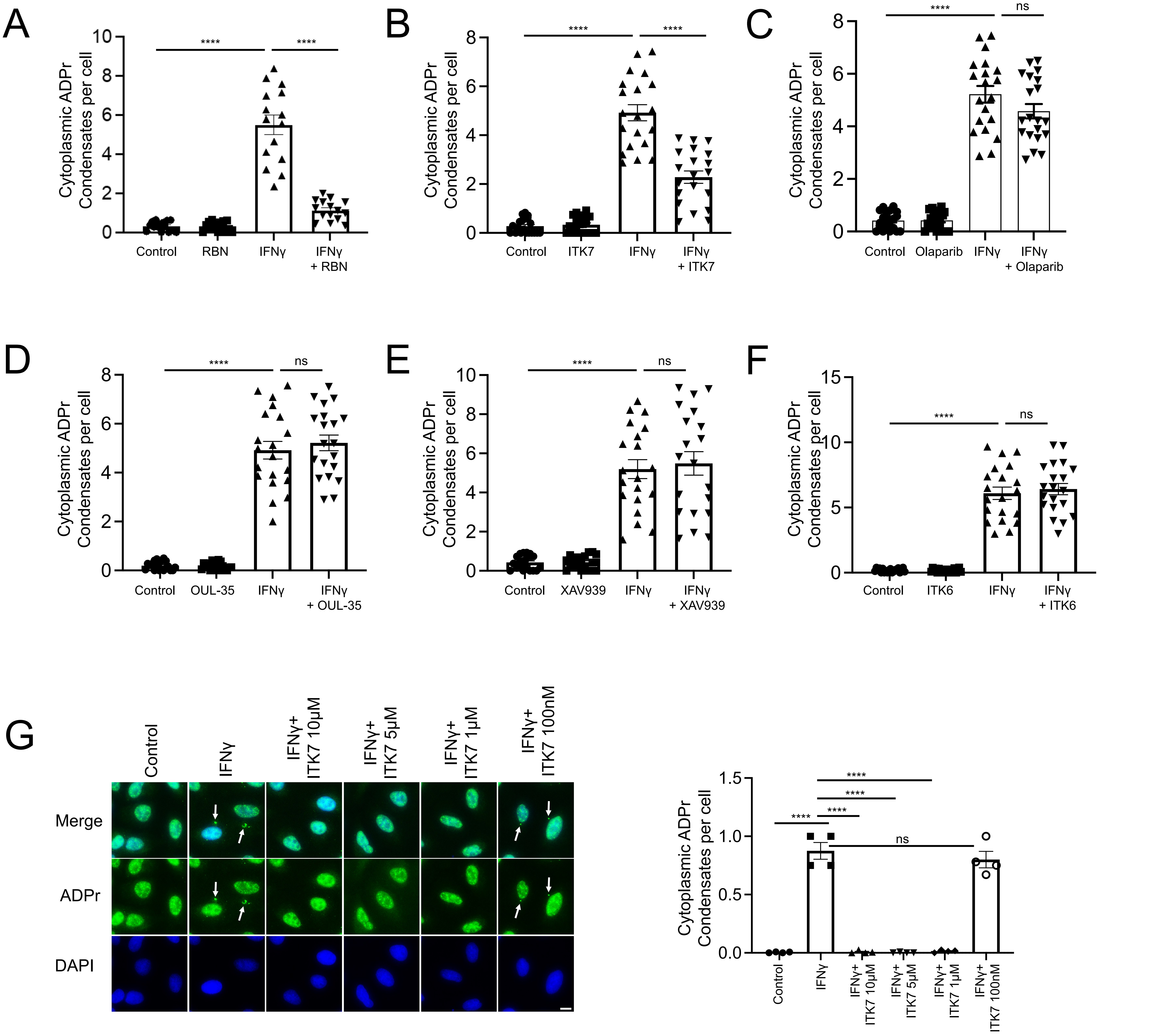


**Figure S2.**

Quantification of ADPr condensates in A549 cells pretreated with different PARP inhibitors for 1 h prior to 24-h IFNγ treatment: (A) 10 µM RBN, (B) 10 µM ITK7, (C) 10 µM Olaparib, (D) 3 µM OUL-35, (E) 10 µM XAV939, and (F) 10 µM ITK6. ****$p < 0.0001$, t-test, ns = not significant, n = 3. (G) ADPr condensate formation was analyzed in cells pretreated with different doses of ITK7 for 1 h prior to 24-h IFNγ treatment. Scale bar, 10 µm

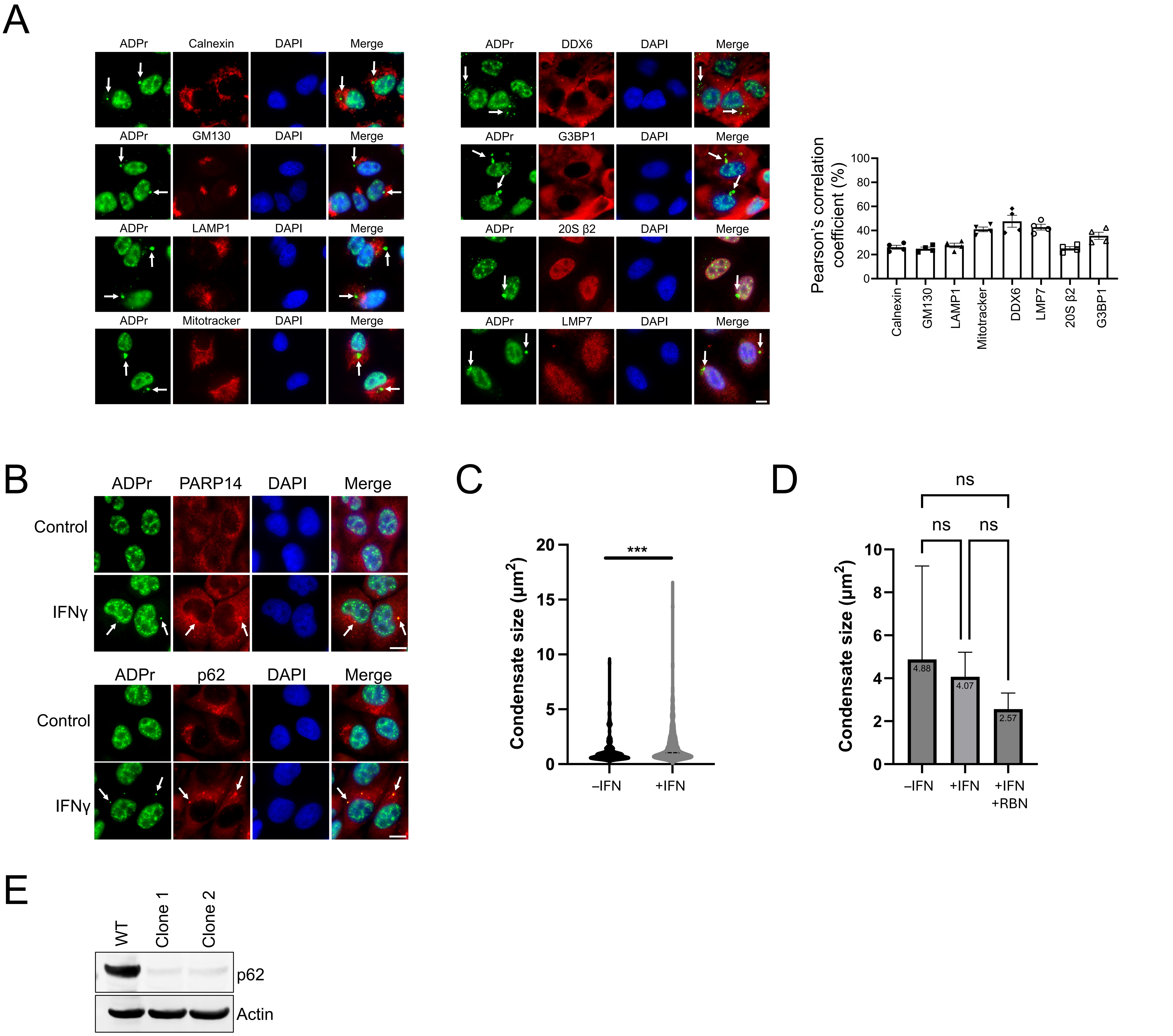


**Figure S3**

(A) A549 cells were treated with IFNγ for 24 h and stained for colocalization of different markers of cellular structures (red) and ADPr (green). The colocalization of ADPr condensates with the cellular structures was calculated by ImageJ software using Pearson's correlation coefficient. (B) A375 cells were treated with IFNγ for 24 h and analyzed for colocalization of ADPr with PARP14 (upper) or p62 (lower). (C) Size quantification of p62 bodies in A549 cells treated with or without IFNγ for 24 h, plotted as a violin plot. ***$p < 0.001$, t-test, n = 182 (–IFN γ) versus 467 (+IFNγ) from three different fields in each group. (D) Size comparison of p62 bodies in FRAP analyses under different conditions for Figure 3C and 4F. ns = not significant, one-way ANOVA. (E) p62 protein levels were measured in shp62 knockdown A549 cell clones generated by lentiviral transduction. Scale bar, 10 μm.

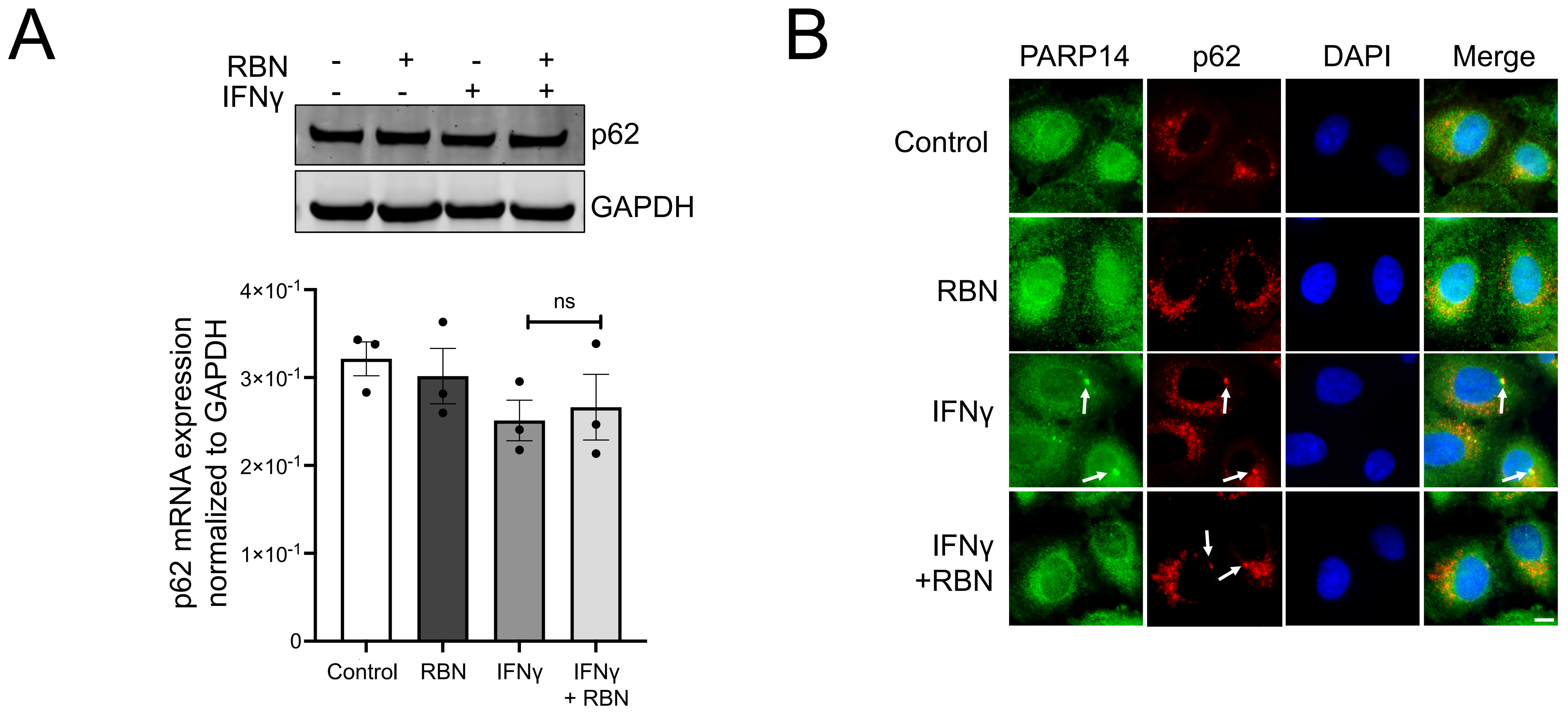


**Figure S4**

(A) p62 protein levels were measured in A549 cells treated with different combinations of RBN and IFNγ, as in Fig 2E. Corresponding mRNA levels of p62, either mock-treated or treated with RBN, IFNγ, or both, were quantitated by qPCR. ns = not significant, t-test, n = 3. (B) The effect of RBN on PARP14 (Abcam) and p62 colocalization was determined in cells pretreated with RBN for 1 h, followed by IFNγ treatment for 24 h. Scale bar, 10 μm.

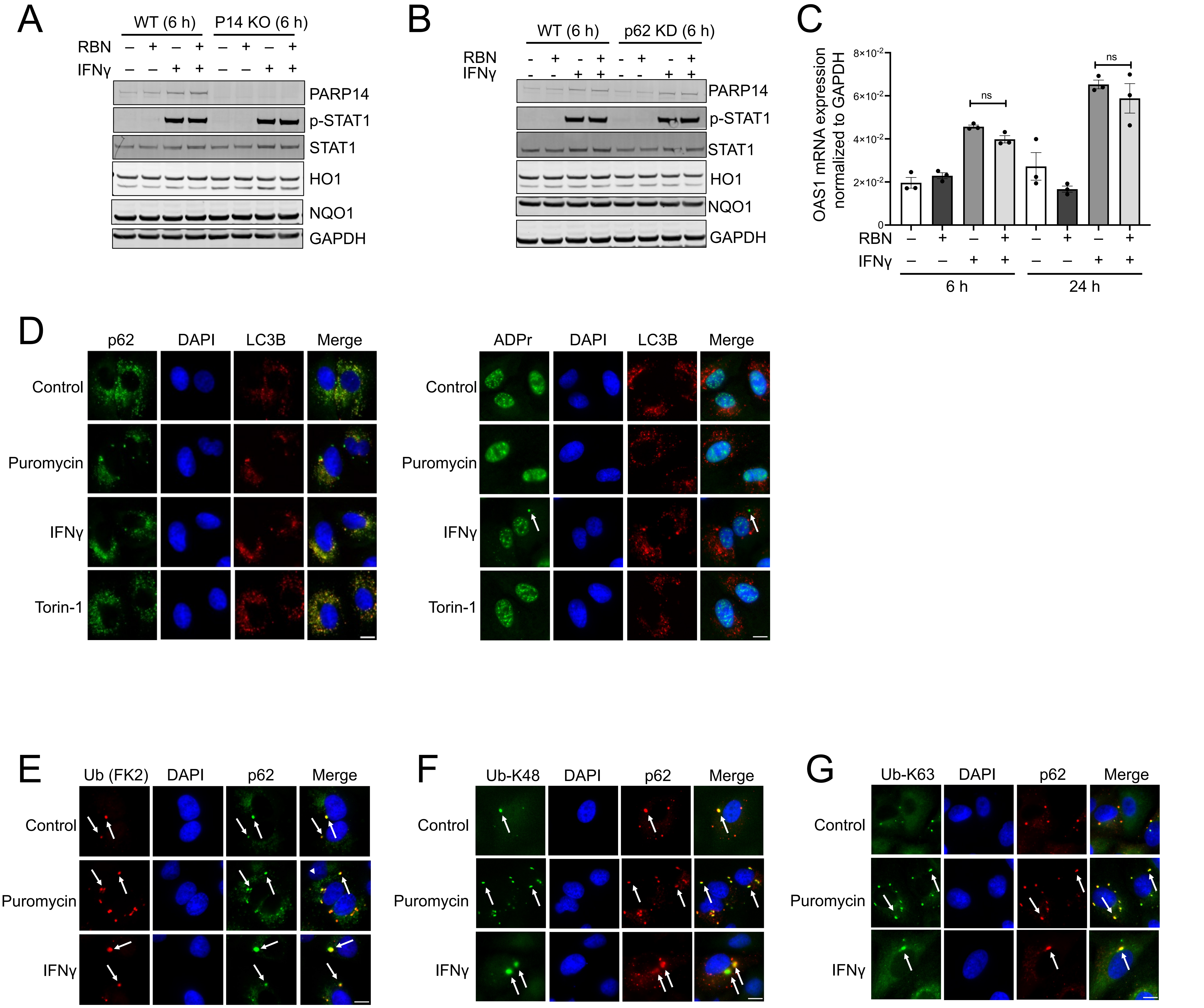


**Figure S5**

(A) Western blot analyses of p-STAT1, STAT1, HO1, and NQO1 in A549 wild-type (WT) and PARP14 knockout (KO) cells treated with different combinations of 1-h RBN pre-treatment and 6-h IFNγ treatment. (B) Western blot analyses of p-STAT1, STAT1, HO1, and NQO1 in wild-type (WT) and p62 knockdown (KD) cells treated with different combinations of 1-h RBN pre-treatment and 6-h IFNγ treatment. (C) qPCR analyses of OAS1 mRNA levels in cells treated with different combinations of RBN pretreatment for 1 h followed by IFNγ treatment for either 6 h or 24 h. ns = not significant, t-test, n = 4. (D) Colocalization of LC3B with p62 or ADPr was assessed in cells either treated with puromycin, IFNγ or Torin-1. (E-G) Colocalization of p62 with (F) Ubiquitin, (G) Ub-K48, (H) Ub-K63 was assessed in cells treated with or without IFNγ or puromycin. Scale bar, 10 μm.

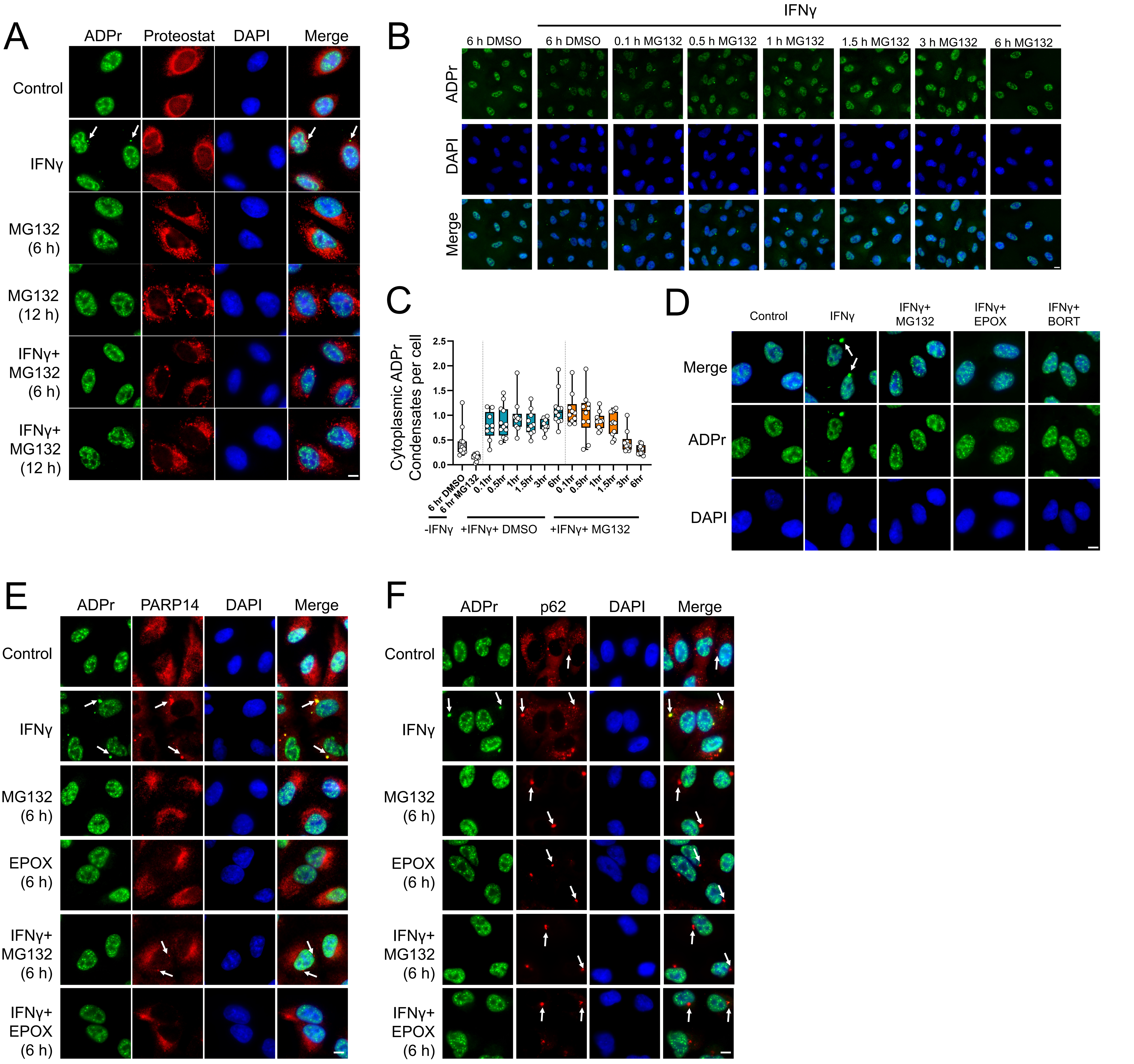


**Figure S6**

(A) Colocalization of ADPr and aggresomes stained with PROTEOSTAT® dye was assessed in A549 cells treated with IFNγ overnight followed by 10 μM MG132 treatment for either 6 h or 12 h. (B) ADPr condensates were monitored after cells treated with IFNγ for 24 h followed by 10 μM MG132 for different time periods as indicated. (C) Quantification of ADPr condensates in IFNγ-treated cells either in the presence of DMSO or 10 μM MG132. (E) Colocalization of ADPr and PARP14 was analyzed in cells treated with IFNγ overnight, followed by the treatment with 10 μM MG132 or 1 μM Epoxomicin for 6 h. (F) Colocalization of ADPr and p62 was analyzed in cells treated with MG132 or Epoxomicin as in panel E. Scale bar, 10 μm
